# Age Distribution, Trends, and Forecasts of Under-5 Mortality in Sub-Saharan Africa

**DOI:** 10.1101/405258

**Authors:** Iván Mejía-Guevara, Wenyun Zuo, Eran Bendavid, Nan Li, Shripad Tuljapurkar

## Abstract

**Background:** Despite the sharp decline in global under-5 deaths since 1990, uneven progress has been achieved across and within countries. In Sub-Saharan Africa, the Millennium Development Goals targets for child mortality were met only by a few countries, and recently new targets were set in goals for Sustainable Development that include the eradication of preventable deaths by reducing neonatal and under-5 mortality rates to at least as low 12 and 25 per 1000 live births by 2030, respectively. As the reduction of preventable deaths has a direct impact on their age distribution, the foci of this study are assessing age patterns, trends over time, and forecasts of mortality rates in Sub-Saharan Africa.

**Methods and findings:** Data came from 104 nationally-representative Demographic and Health Surveys with full birth histories from 31 Sub-Saharan African countries from 1990 to 2016 (a total of 448 country-years of data). We assessed the distribution of age at death through the following demographic model. First, we used a direct method for the estimation of death rates with full-birth histories from survey data to construct age profiles of under-5 mortality on a monthly basis. Second, a two-dimensional P-spline approach was used to smooth out raw estimates of death rates by age and time. Third, a variant of the Lee-Carter model, designed for populations with limited data, was used to fit and forecast age profiles of mortality. We used mortality estimates from the United Nations Inter-agency group for Child Mortality Estimation to adjust, validate and minimize the risk of bias in survival, truncation, and recall in mortality estimation.

Our study has three salient findings. First, we observe a monotonous decline of death rates at every age in most countries, but with notable differences in the age-patterns over time. Second, our projections of continued decline of child mortality differ from existing estimates from the United Nations Inter-agency group for Child Mortality Estimation in 5 countries for both neonatal and under-5 mortality. Finally, we predict that only 5 countries (Guinea, Liberia, Rwanda, Tanzania, and Uganda) are on track to achieve the sustainable development goal targets on child mortality by 2030. Poor data quality issues that include bias in the report of births and deaths, or age heaping, remain a limitation of this study.

**Conclusions:** This study is the first to combine full birth history data and mortality estimates from external reliable sources to model age patterns of under-5 mortality across time in Sub-Saharan Africa. We demonstrate that countries with a rapid pace of mortality reduction across ages would be more likely to achieve the sustainable development goal targets of child mortality reduction. Our mortality model predicts that if neonatal and under-5 deaths decline at the rates observed during the last 25 years, only 5 countries would reach those targets by 2030, 15 would achieve them between 2030 and 2050, and 11 afterwards.

## Introduction

Under-5 mortality analysis has been critical in evaluating progress towards the Millennium Development Goal 4 (1), and more recently towards the Sustainable Development Goal 3 (SDG-3), which aims to reduce neonatal mortality to fewer than 12 per 1000 live births and under-5 mortality rates to at least as low 25 per 1000 births by 2030 (2). The monitoring of child survival is conducted by the United Nations Inter-Agency group for Child Mortality Estimation (UN IGME) (1), which has adopted a methodology for child mortality estimation (3,4), and regularly updates child mortality levels and trends around the world (4).

The most recent estimates from UN IGME revealed outstanding progress as the total number of under-5 deaths dropped from 12.6 million in 1990 to 5.6 million in 2016. Yet, this latter figure means over 15,000 deaths every day, and reflects uneven progress among and within countries, particularly in Southern Asia and Sub-Saharan Africa (SSA) (4–6), where about 80 per cent of under-5 deaths occur (4). In particular, despite the impressive reduction of 57% of under-5 deaths in SSA between 1990 and 2016 (Average Annual Rate of Reduction–ARR = 3.2), it remains the region with the highest under-5 mortality in the world (4).

Recent evidence reveals uneven trends in the reduction of child mortality rates in low- and middle-income countries (LMICs), particularly for specific population subgroups: by sex (7,8); by wealth status–with absolute disparities in mortality declining between the poorest and richest households but with persistent relative differences (9,10); over space, with substantial spatial heterogeneity within countries (11) but some convergence at subnational levels (12); and for some causes of death (6,13). Methodological work has addressed the inadequacy of traditional life table models applied to child mortality in SSA (14), and small area smoothing with data from sample surveys and demographic surveillance systems (15). But little attention has been paid to the age distribution of deaths, although recent studies (5,16,17) do report differences in some causes of death between neonates and older children.

It is well documented that the reduction in child mortality was a key factor for the change in the age distribution of mortality and the increase of life expectancy experienced in the developed world during the XX Century (18,19), as life expectancy is particularly sensitive to mortality reductions at younger ages (20). With subsequent declines in child mortality over time, we expect that infant deaths in SSA countries will tend to concentrate in the first month of life as post-neonatal conditions improve due to the eradication of exogenous mortality causes, and then endogenous causes-those that may not yield medical progress, would persist (21,22). This change relates to the classical epidemiological transition model (23), which states that childhood survival (particularly at ages 1-4) benefits the most from the shift of disease patterns and the increase in life expectancy as infectious diseases are progressively displaced by ‘degenerative and man-made diseases’; although the duration, pace, timing, and determinants have been subject to criticism (24–26)

The main purpose of this study is three-fold: 1) contribute to filling the gap in modeling fine-grained dating mortality patterns for under-5 children, 2) the analysis of trends in the age at death distribution for under-5 children, and 3) the forecasting of age patterns and mortality levels by country in SSA. Our analytical approach provides important insights for the evaluation and assessment of the millennium and sustainable targets of child mortality; the systematic evaluation of data errors in sample surveys and its impact on age patterns of mortality; and the potential for identifying a wide range of epidemiological situations and trajectories.

## Methods

### Data Sources

Data are birth histories from 104 Demographic and Health Surveys (DHS) from 31 SSA countries from the period between 1990 and 2015. The DHS program collects health and demographic information mostly for women in reproductive age (15-49 years old) and their children, commonly based on a two-stage stratified cluster sampling design that defines strata by region and by rural-urban within each region. In the first stage, primary sampling units are selected randomly with probability proportional to sampling size from a list of census enumeration units or tracks that cover the entire country and defined according to the most recent sampling frame, usually the latest census when available. In the second stage, DHS survey personnel select households systematically from a list of previously enumerated households (27).

### Full birth histories and retrospective mortality data

Full birth history (FBH) data are available for individual women in DHS surveys, including up to 20 previous births for every eligible woman–usually women 15-49, where the respondent mother is asked about the date of birth of each of her ever-born children, and the age at death if the child has already died (Hill 2013).

FBH data permits the estimation of death rates for up to 25 years before the survey (Hill 2013). We used retrospective information from FBH data, following the statistical guidelines from Pedersen and Liu (29). We selected the time periods recommended by those authors for the estimation of infant mortality rates and for the countries and survey-years considered in their study that matched our sample; and for subsequent survey years that were available after their publication, we considered the time interval used in the latest survey included in that study or a 5-year period for countries that were not included. For each country, we then estimated neonatal (NMR), infant (IMR), and under-5 (U5MR) mortality rates retrospectively, starting with year 1990 (or later, for some countries) to focus on the period 1990 to 2015.

### Demographic Methods

We built our demographic model as follows: 1) we computed conditional life-table age distribution of under-5 deaths from survey data, 2) we adjusted our mortality profiles to match neonatal, infant, and under-5 mortality rates from UN IGME estimates, and 3) we smooth-out the age mortality profiles, and fit and forecast them using a modified version of the Lee-Carter model. Details for each step are described in what follows.

### 1) ‘Conditional’ life-table age distribution *of* under-5 deaths

We constructed life-table age distributions of death, based on estimated death rates obtained from death reports by households, and birth history data from DHS (28). We assigned deaths and exposure time across each calendar year on a monthly basis. Estimates of age-specific death rates *m_[x]_* ([x] stands for age in months) considered the contributions of children in the survey to the number of events and total time to event (28). We computed period life-table probabilities of dying, *q_[x]_* (probability of dying between month *x* and month *x*+*1*), assuming that deaths are distributed uniformly across every single month age range,

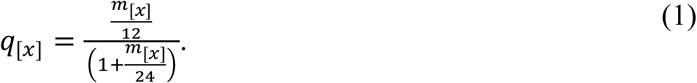

We estimated NMR, IMR, and U5MR rates using this methodology and the formulae (28),

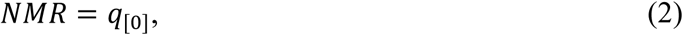

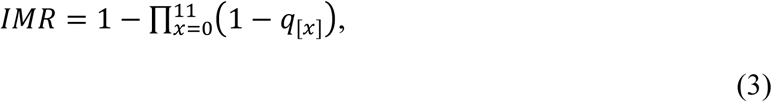

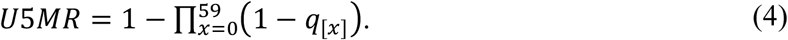

### 2) *UN* IGME neonatal, infant and under-5 mortality adjustment

Direct estimates of under-5 deaths based on FBH are prone to measurement errors as the information is reported directly by living mothers (survivor bias) or due to an upper age limit that is usually considered as an eligibility criterion for surveyed women (truncation bias: eligible women are usually in the age range 15-49) (28,30). Survivor bias is particularly relevant in countries with extended periods of high HIV prevalence (31).

Because the lack of high quality vital registration systems for the countries in our sample, we used mortality estimates from the UN IGME group (1,4) that are designed to mitigate bias and error (3), to validate and adjust our mortality estimates. Specifically, a) we adjusted our raw monthly death rates to match the UN IGME estimates for the neonatal (<1m), post neonatal (1-11m), and childhood periods (12-59m); and b) we used the UN IGME rates to validate the age-adjusted trajectories obtained after smoothing and fitting our model (more details in the following section). For (a), the monthly probabilities of surviving in equations (2)-(4) *p*_[*x*]_= [l — *q*_[*x*]_]were adjusted proportionally to match UN IGME estimates of NMR, IMR, and U5MR exactly, resulting in three measurement errors *d*_*M*_=1-f_*M*_ (M=nmr, pmr, cmr), where *f*_*M*_ stands for the adjustment factor applied respectively to *[1]p[o]*, *[11]p[1]*, and *[59]p[12]* (*[a]p[x]* is the probability of surviving from the age month *x* to <u>*x*+*a*</u>).

Our direct unadjusted estimates of neonatal mortality differ by as much as 2% from the UN IGME values, in contrast with the post neonatal and child periods that were highly concordant (i.e., with practically no adjustment required) for the majority of year periods. Our unadjusted estimates of neonatal rates were noisy over time, as we expected from our use of retrospective data, and the noise would have been greatly reduced for many countries by a moving average (no average is used in the analyses reported here). In **S1 Fig.**, we show the magnitude of the error (*d_*M*_*) after the adjustment we made to neonatal rates (*f_nmr_*), and the much smaller adjustments we made to post-neonatal (*f_pmr_*) (period between ages 1 and 11 months), and child mortality (*f_cmr_*) rates (between ages 12 and 59 months) to fit exactly the UN IGME estimates.

### 3) Fit and forecasting of mortality trajectories

We used a two-dimensional P-Spline smoothing and generalized linear model to smooth our calibrated mortality profiles over ages and years, assuming that the number of deaths at a given rate are Poisson-distributed (32). A variant of the Lee-Carter (LC) (33) model (Li-Lee-Tuljapurkar–LLT) is applied to the age mortality profiles after smoothing. In contrast with the standard LC model (34), the modified version is suitable for mortality profiles using datasets that contain multi-year gaps and provides various measurement errors (95% unbiased, narrow, and wide error bounds) if data are available at least for 3 periods. This characteristic is particularly relevant for this study because our DHS life histories contain year gaps for some countries even after we augmented the periods of analysis using the retrospective information when trying to fill those gaps. This augmentation did allow us to overcome the requirement of the 3 years of data in the modified LC approach (33). For age [x] and year, *t* the Lee-Carter model that we fit has the form,

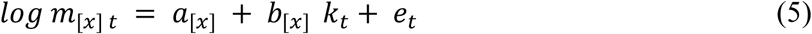

where the first two terms on the right are estimated in a singular-value decomposition step, and the last term is an error term whose variance is estimated as described by Li et al. (33). We measured the goodness-of-fit of the LC model as the percentage of the variance explained of the mortality profile (*m*_[*x*]*t*_–after the adjustment to match UN IGME estimates) by the first principal component of the singular-value decomposition (details in **S1 Text**) (35). For most countries, the LC model captured more than 90% of the total variation of smoothed under-5 mortality, from 92% in Angola to 98% in Nigeria; except for Niger, where it accounted for 81% of the total (**S1 Table**). The resulting fits were used to generate smoothed point estimates (the median Lee-Carter values) of age-specific death rates within the 1990-2016 period, and NMR, IMR, and U5MR mortality rates. These estimated rates fell within the credible intervals reported in the latest revision of the UN IGME model (see **Figure 1**). The dots in that figure represent mortality estimates from our LC model and the shaded areas credible intervals reported by UN IGME. Around 97%, 85%, and 88% of our neonatal, infant, and under-5 mortality estimates fell within those credible limits, respectively (we dropped Rwanda estimates from the period between 1990-1993 as the resulted mortality rates looked unrealistically high).

**Fig 1.**
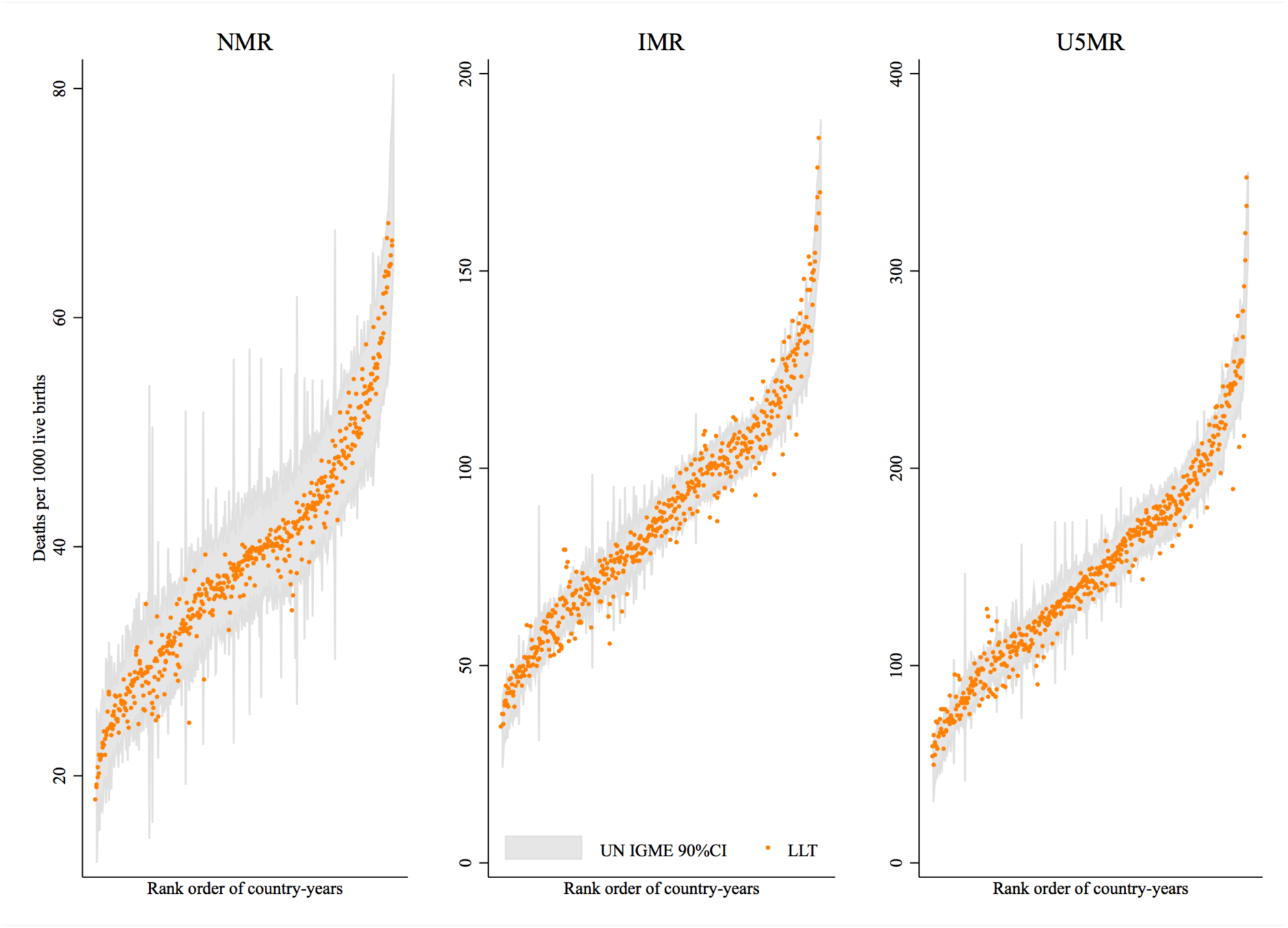
Neonatal (NMR), infant (IMR), and under-5 (U5MR) mortality rates estimated using the fitted age profiles from the Li-Lee-Tuljapurkar (LLT) model and 90% credible intervals (CI) from UN IGME estimates for selected year time periods (between 1990 and 2016) in 31 Sub-Saharan African countries. Country-years represented in the X-axis were sorted on the basis of mortality levels. We retrieved UN IGME estimates from (IGME 2017).

## Results

### Trends and forecasts of age-specific mortality patterns

We used our LLT model to fit and forecast under-5 death rates for every country. The forecasts of trajectories were for the years 2030 and 2050 (predictions for Lesotho were precluded by the poor quality of data and great uncertainty in the estimates and uncertainties). Most countries display a monotonous decline of death rates at every age, but there are notable differences in the age-patterns over time. Death rates by age for all countries are in **Fig. 2**, including fitted (1990 and 2015) and predicted (2030) age-specific mortality trajectories. For instance, Chad, Nigeria, Rwanda, and the United Republic of Tanzania (Tanzania hereafter) are representative countries with different trajectories of mortality reduction. Chad and Nigeria are countries with low ARR (below 3.0–the median ARR of the countries in our sample), whereas Rwanda and Tanzania represent countries with high ARR (above 3.0). For Chad, mortality fell most rapidly at ages between 1 and 3 years, leading to a decidedly uneven pattern (by age) in 2015 and persisting to 2030. However, in Nigeria, although death rates during the second year of life decreased more rapidly in the initial period, mortality at all ages eventually declined at similar proportional speeds and the mortality profile remains linear with age to 2030. Our prediction model for both countries indicates an improvement in the mortality levels at all ages by 2030, but with more uncertainty in Chad. Mortality patterns in Rwanda and Tanzania are similar, the transition from high to low mortality across the under-5 period starts with a sharp decline in infant (2-11m) and child (12-59m) mortality, followed by a less rapid decline in neonatal (<1m). In both countries, though, the mortality profile is steepest during the infant compared to the child period across time. Our model also predicts significant declines of mortality by 2030 at all ages, the mortality curve in Rwanda becoming increasingly rectangular, i.e. concentrating near birth and then falling sharply and rapidly with age.

**Fig 2.**
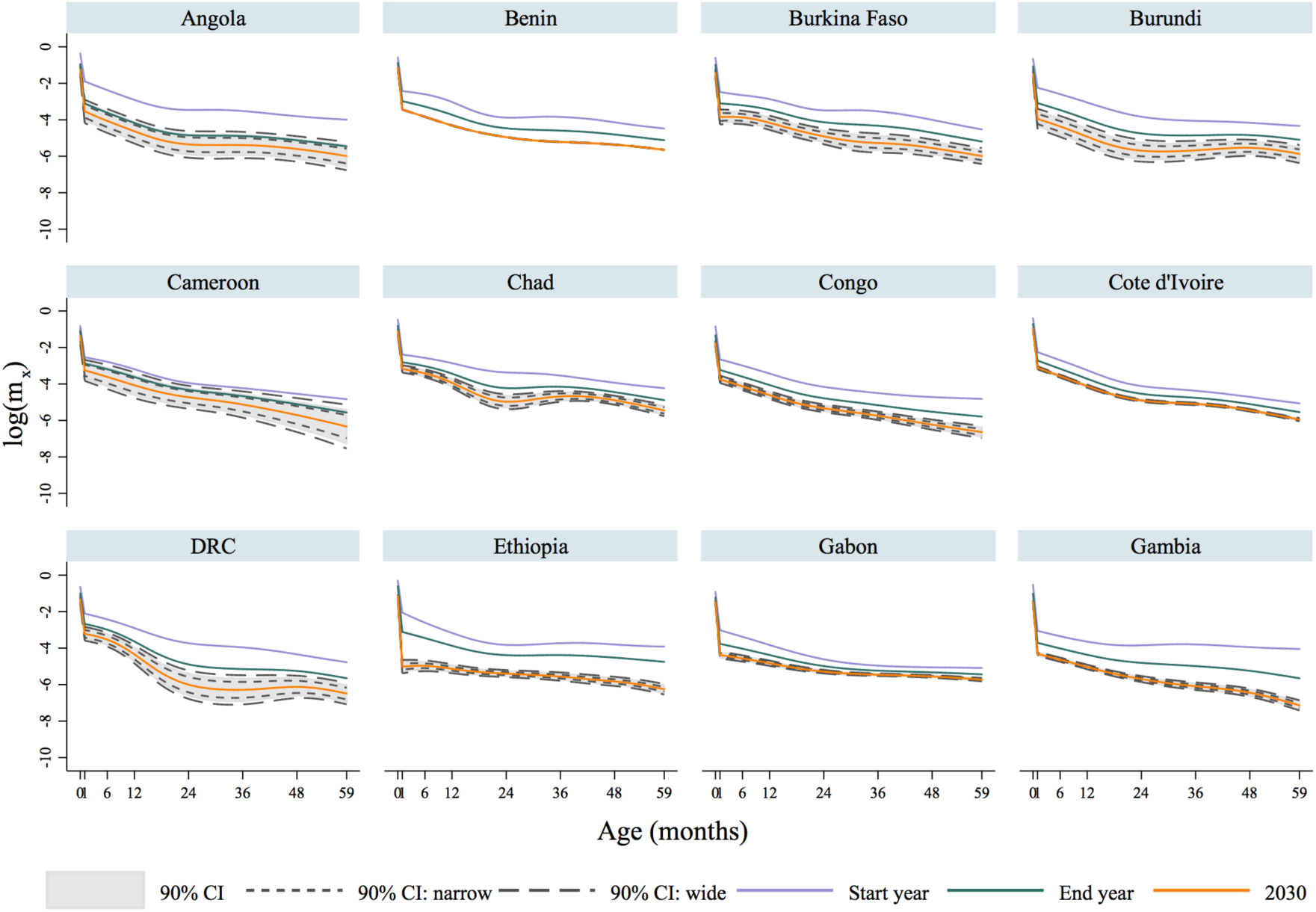
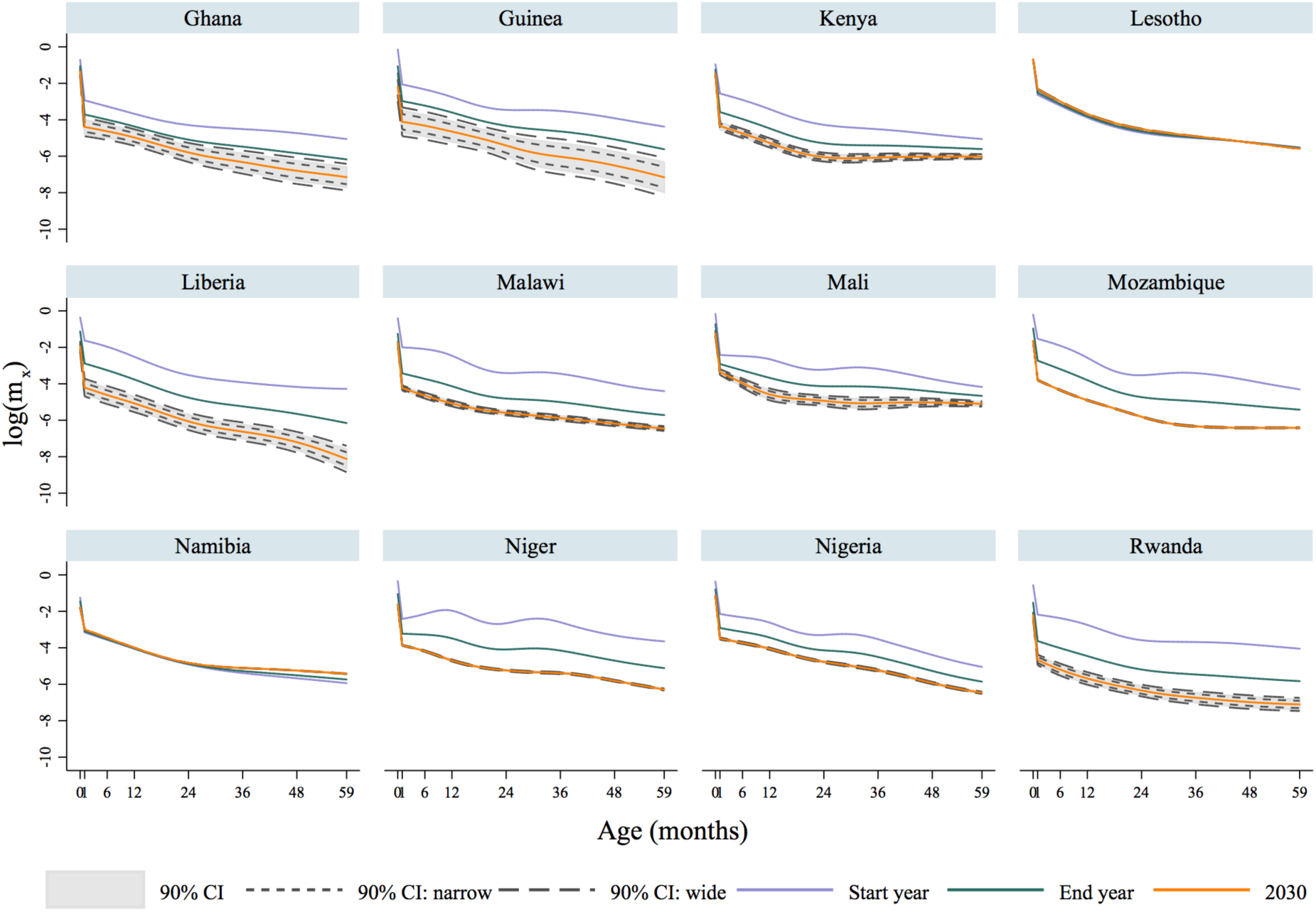
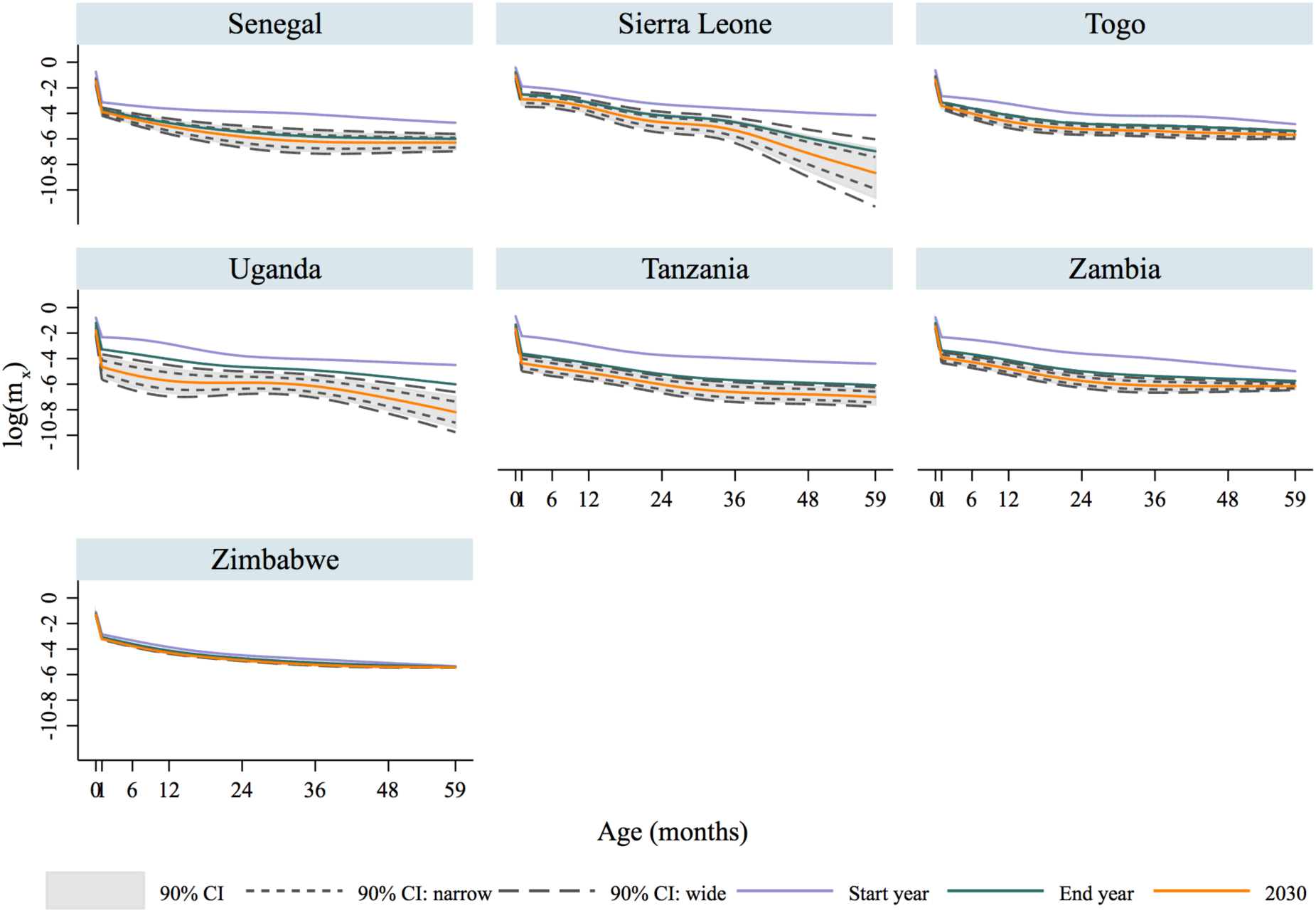
Lee-Carter fit and prediction of age patterns of death rates for under-5 children in 31 Sub-Saharan African countries. Source: Author’s estimates using the Li-Lee-Tuljapurkar model (Li, Lee, and Tuljapurkar 2004) with data from the DHS program.

Our forecasting model revealed that countries that experienced a low pace of under-5 mortality decline in the past would most likely fall short in achieving the SDG-3 by 2030, in contrast with those experiencing more substantial or accelerated reductions. For example, the ARR (the formula for its estimation is in **S1 Text**) for neonatal and under-5 mortality in Nigeria between 1991 and 2013 of 1.5 and 2.6, respectively, would remain the same by 2030 and 2050, and that would prevent the country for achieving the SDG-3 targets-neither in 2030 nor in 2050. In contrast, the ARR of neonatal and under-5 mortality in Guinea would increase from 3.6 to 4.7 by 2030 and, as a consequence, the country would likely reach those targets (**S1 Table** includes predicted ARRs for all countries). Rwanda and Tanzania, two countries that have achieved the MDG-4 (36), would likely meet the SDGs of neonatal and under-5 mortality reduction by 2030, according to our predictions. In summary, we predict that only 5 countries are on track to achieve the SDG-3 by 2030 (Guinea, Liberia, Rwanda, Tanzania, and Uganda) (**Figure 3**), and if the observed pace of mortality reduction continues, and considering the uncertainty predicted by our model (based on our estimated unbiased error bounds), we predict that only 15 additional countries would achieve the goal between 2030 and 2050 (Angola, Burkina Faso, Burundi, Cameroon, Congo, Gabon, Gambia, Malawi, Mozambique, Namibia, Niger, Senegal, Sierra Leone, Togo, and Zambia), and 11 more would make it after 2050 (Benin, Chad, Cote d’Ivoire, Democratic Republic of the Congo-DRC, Ethiopia, Ghana, Kenya, Lesotho, Mali, Nigeria, and Zimbabwe) (**Fig. 3**).

**Fig 3.**
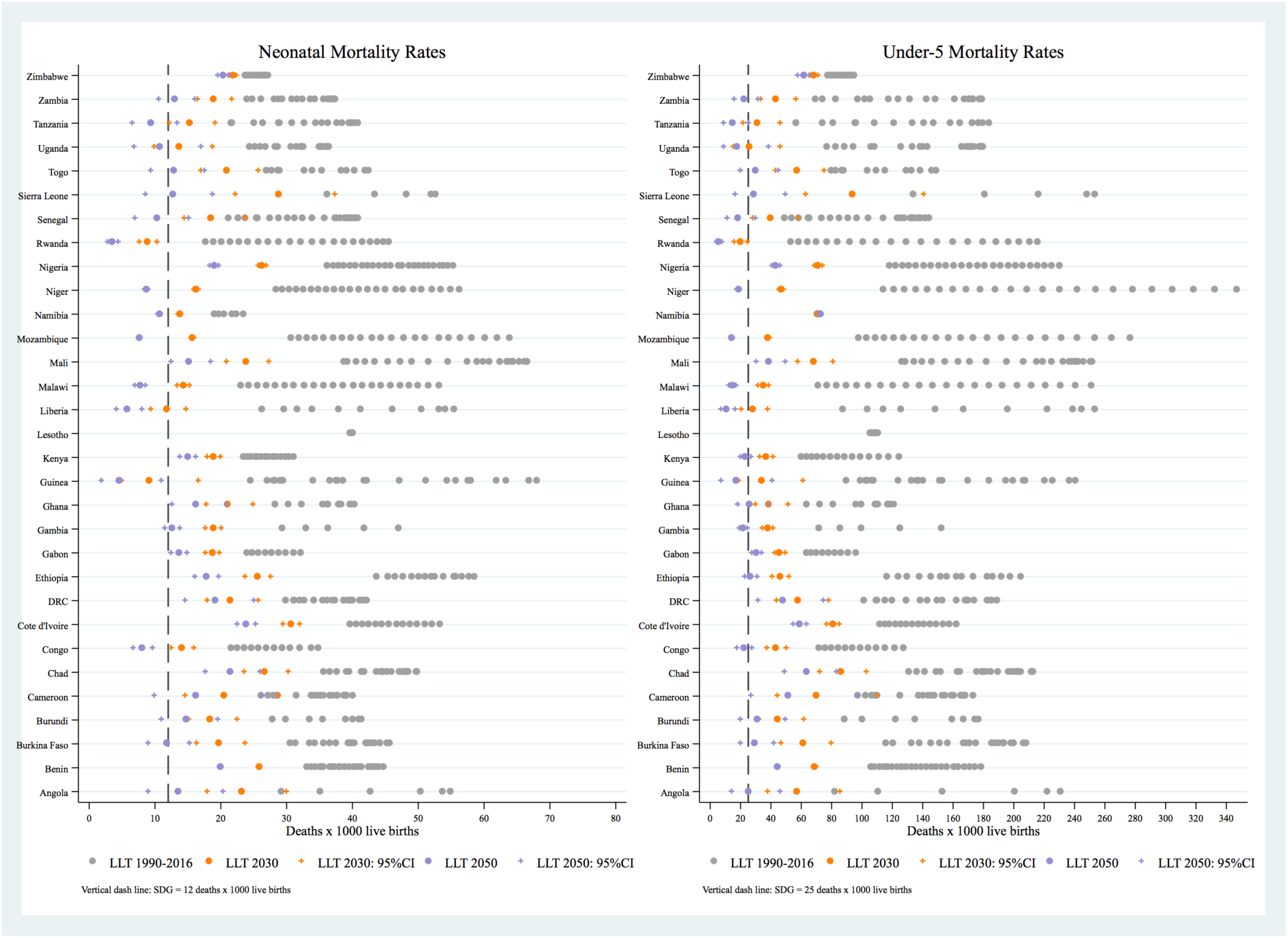
Forecasts of neonatal and under-5 mortality by 2030 and 2050 based on the LLT model, and assessment of SDG-3 targets in 31 Sub-Saharan African countries. Predictions for Lesotho were precluded by the poor quality of data and great uncertainty in the estimates and uncertainties. We report unbiased error bounds for our prediction models for 2030 and 2050.

Our assessment of the SDG-3 progress is line with estimates from UN IGME in the majority of countries. We identified 5 countries with discrepancies in both neonatal and under-5 mortality predictions: Guinea, Liberia, Senegal, Togo, and Tanzania. In Guinea, Liberia, and Tanzania the difference is that we predict an early transition towards the SDG-3 targets (**S2 Fig.**). Further comparisons with predictions from the Institute for Health Metrics and Evaluation (IHME) revealed that our estimates (and those from UN IGME) are less optimistic in general, although we found an overlap in 23 and 20 countries (out of 31) for neonatal and under-5 estimates, respectively (i.e., IHME predictions are within our estimated error bounds or ours within IHME error bounds) (**S3 Fig.**).

## Discussion

This paper describes three novel findings. First, we advanced the modeling of age patterns of under-5 mortality for detailed age groups using FBH data from Sub-Saharan Africa, providing important information on under-5 mortality patterns, and representing a step forward in the analysis of changes in age patterns of mortality across time and by country. We used the latest refinements in the estimation of child mortality based on full-birth histories from survey data, and adjust and validate our rates using official estimates of under-5 mortality rates that are derived from a robust model (1,3). Second, we made probabilistic projections of age patterns of mortality by 2030 (and where possible to 2050) in order to assess progress towards the SDGs of child mortality reduction. In making that assessment, our use of probabilistic methods allowed us to account for different degrees of uncertainty. Our predictions are consistent with estimates from UN IGME as we found discrepancies for only 5 countries in the timing where the SDG-3 would be achieved. Third, we predict that Guinea, Liberia, Rwanda, Tanzania, and Uganda are on track to achieve the SDG-3 for child mortality reduction, and 11 countries would achieve them only after 2050.

This study is in line with previous findings of under-5 mortality reduction in SSA (4), but goes further by showing the reductions in age-specific death rates in a monthly basis for the majority of countries in the region. It also identified heterogeneities in the trends and age patterns of mortality decline across countries, with important lags in most countries that would prevent them to reach the SDGs targets. Other countries, like Rwanda and Tanzania, have succeeded in achieving the Millennium Development Goals 4 for child mortality reduction and would likely meet the SDG-3.

However, despite the accelerated progress in the reduction of under-5 deaths observed in some countries, the mechanisms and underlying factors leading to improvements in child survival require further examination. The global mortality rank in pneumonia and diarrhea deaths in under-5 children, the two diseases responsible for about 25% of all the deaths that occurred in under-5 children in 2015, reveals that 72% of the global burden of pneumonia and diarrhea child deaths occurred in just 15 countries (38). Seven countries in our study sample (Angola, Chad, Democratic Republic of Congo-DRC, Ethiopia, Niger, Nigeria, and Tanzania) are among them, but only Chad and Nigeria have registered ARR below 3.0% (**S1 Table**). That is, countries that have achieved significant progress in the reduction of mortality in the past (i.e., Ethiopia, Niger, and Tanzania) still face significant challenges. In addition, although previous studies did not attribute the emergence of HIV as the leading cause of modifying preexisting patterns of under-5 mortality in some SSA regions (14), our findings indicate that Lesotho, Namibia, Malawi, Zambia, and Zimbabwe (countries largely affected by the HIV/AIDS epidemy) showed signs of poor data quality, unreliable fit and predictions (Lesotho, Namibia, and Zimbabwe), but also substantive progress in child mortality reduction during the previous two decades for some of them (Malawi and Zambia).

Recent studies for countries that achieved the MDG-4 targets revealed important insights into the determinants of change, coverage, intervention and implementation of policies that have succeeded in specific contexts. Ethiopia have developed multisectoral policy platforms that integrates child survival and specific health goals within macro-level policies and programmes (39). Niger has developed policies intended to increase access to child health services, the use of mass campaigns and programming for nutrition (40). Tanzania put high political priority to child survival, with consistent increases in funding and focused on the implementation of high-impact interventions at lower levels of the health system, although to the detriment of mothers and neonates (41). After the Rwandan genocide in 1994 that led to the death of more than 1 million people and the devastation of the health system (42), the country embarked on ambitious programmes to provide equitable health services that resulted in the improvement of health equity and child survival. The rebuilding of the health system included notions of ready access, accountability, and solidarity, as well as the implementation and scale-up of community-based health insurance and performance-based financing systems (43).

Our study builds on previous research that examines patterns of mortality as a way to understanding the sources of error or true epidemiological patterns that are not captured by model life table approaches for SSA (14,15,44–46), and stressing the importance of using new methodological approaches and complementary sources of data (15). In addition, we complement previous studies that analyze trends and prediction of mortality rates worldwide (4–6,16).

To the best of our knowledge, our study is the first to construct under-5 mortality patterns from narrow-age groups using a LLT model for the assessment of trends and prediction of under-5 mortality in SSA with uncertainty. Specifically, we made predictions of mortality rates by 2030 for the assessment of the SDG-3 targets, and by 2050 to evaluate which countries in our predictions would meet the SDG-3 targets by then if they fall short to do so by 2030. Our mortality patterns provided evidence of an acceleration of mortality decline and substantive changes in age mortality patterns in countries with higher rates of child survival. In particular, we observed in certain countries that the distribution of deaths would follow a pattern that is becoming increasingly rectangular, having an increasingly flat down and sharp upslope. We refer to this phenomenon as the “early rectangularization” of the under-5 mortality curve, a phenomenon similar but very distinct in nature from the hypothesized rectangularization at old ages, extensively studied in the literature for older populations (47–49). Further analyses of compression and convergence of early mortality would provide more insights about this phenomenon.

The findings of this study should be judge in light of several limitations. First, the gold standard for the analysis of mortality in more developed countries relies on the existence of high quality vital registration systems, but those systems are inexistent or deficient in the countries included in our study (50). Second, this study relies on self-reported information from life histories available in nationally-representative surveys, which are subject to several sources of error. Third, because of the nature of the survey data, we are not able to make a detailed assessment of the underlying causes of mortality reduction across the under-5 period.

## Conclusion

This study contributes to the development of detailed age patterns of mortality for under-5 children, and stresses their importance in the monitoring of child survival of specific age groups to identify distinct patterns of mortality decline at early ages in most countries of SSA. Our estimates and forecasts relied on a robust LLT model that was suitable for our data with year gaps, providing different degrees of uncertainty, and capturing most of the variation of under-5 mortality in the SSA region. Its accuracy could be refined if further reliable sources of information become available, such as the development of new vital registration systems. It should also be considered in the design and scale-up of targeted interventions intended to accelerate progress towards achieving the SDG-3 targets for child mortality reduction. Future research should explore a detailed assessment of age inequality in early mortality, compression and convergence, as well as the true relationships between age patterns of mortality and epidemiological trajectories.

## SI Supporting Information

**SI Fig.**
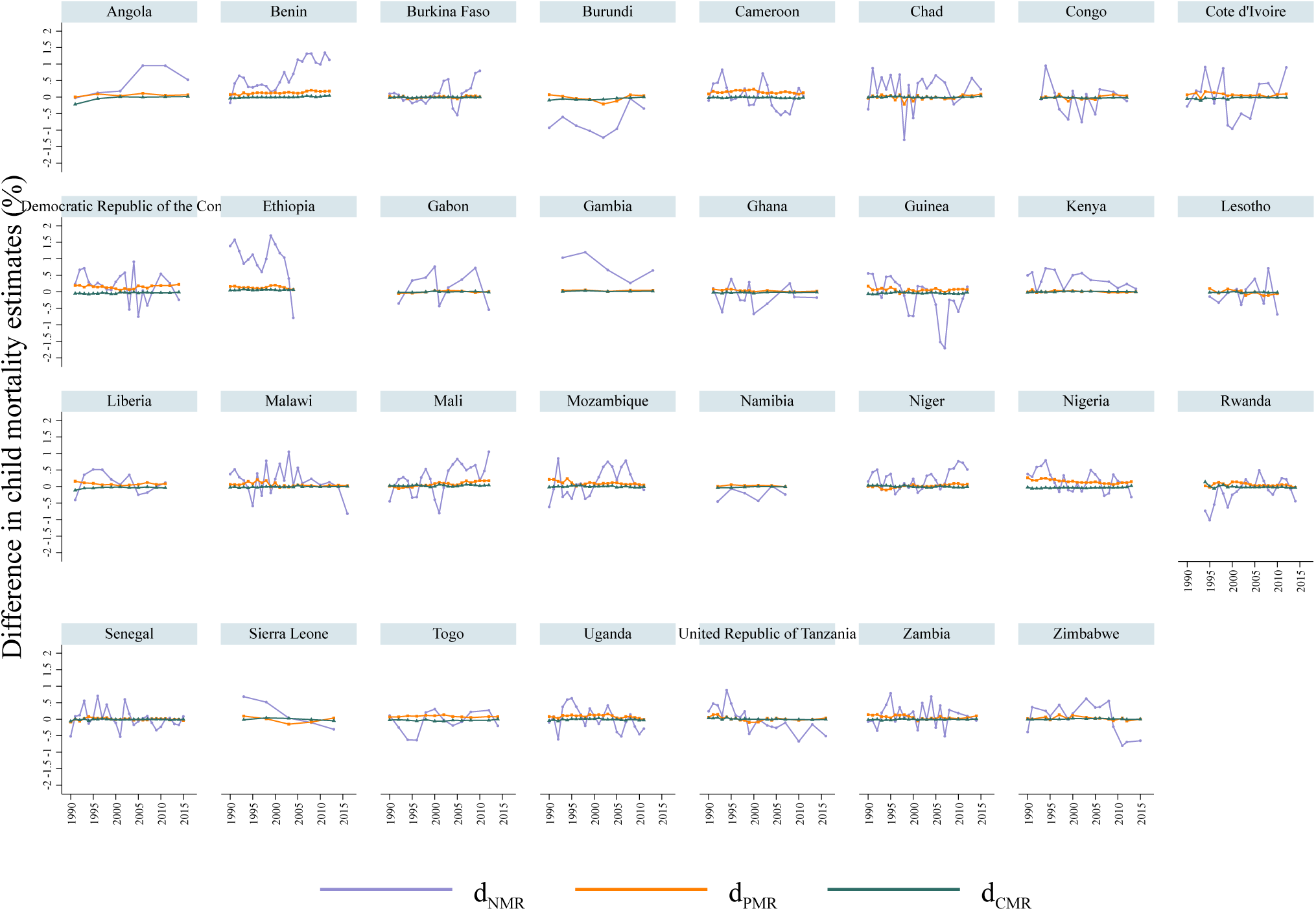
Difference in survival rates that resulted after adjusting DHS data to match United Nations Inter-agency Group for Child Mortality Estimation (UN IGME) estimates for the neonatal (*d*_*NMR*_), post neonatal (d_*PMR*_), and child (*d*_*CMR*_) period from 31 Sub-Saharan African countries by year **Source:** Authors’ estimates using data from the DHS Program.

**S2 Fig.**
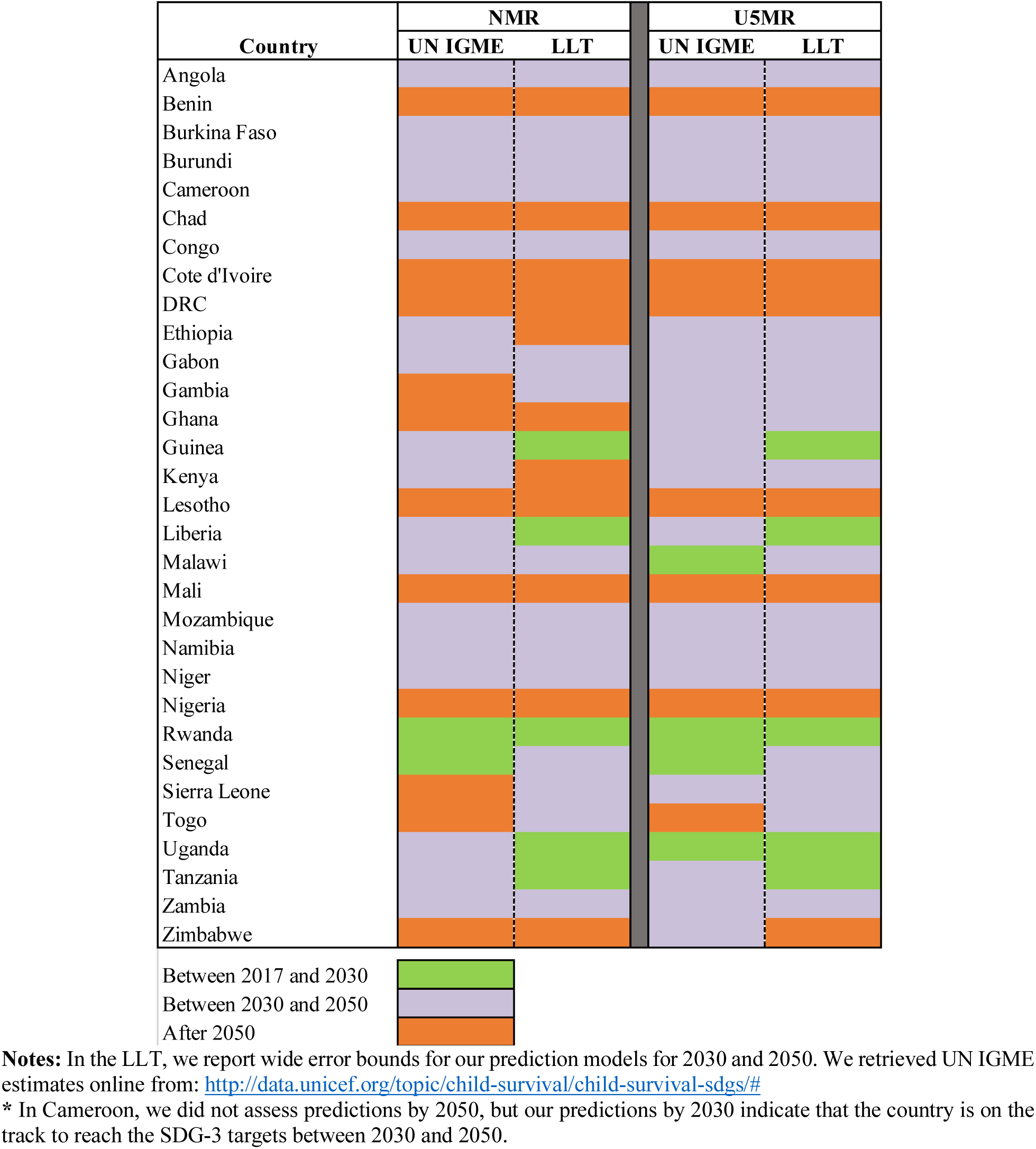
Assessment of the Sustainable Development Goal-3 targets for neonatal and under-5 mortality rates by 2030 and 2050 based on estimates from the Li-Lee-Tuljapurkar (LLT) model and from United Nations Inter-agency Group for Child Mortality Estimation (UN IGME) for 31 countries from Sub-Sharan Africa.

**S3 Fig.**
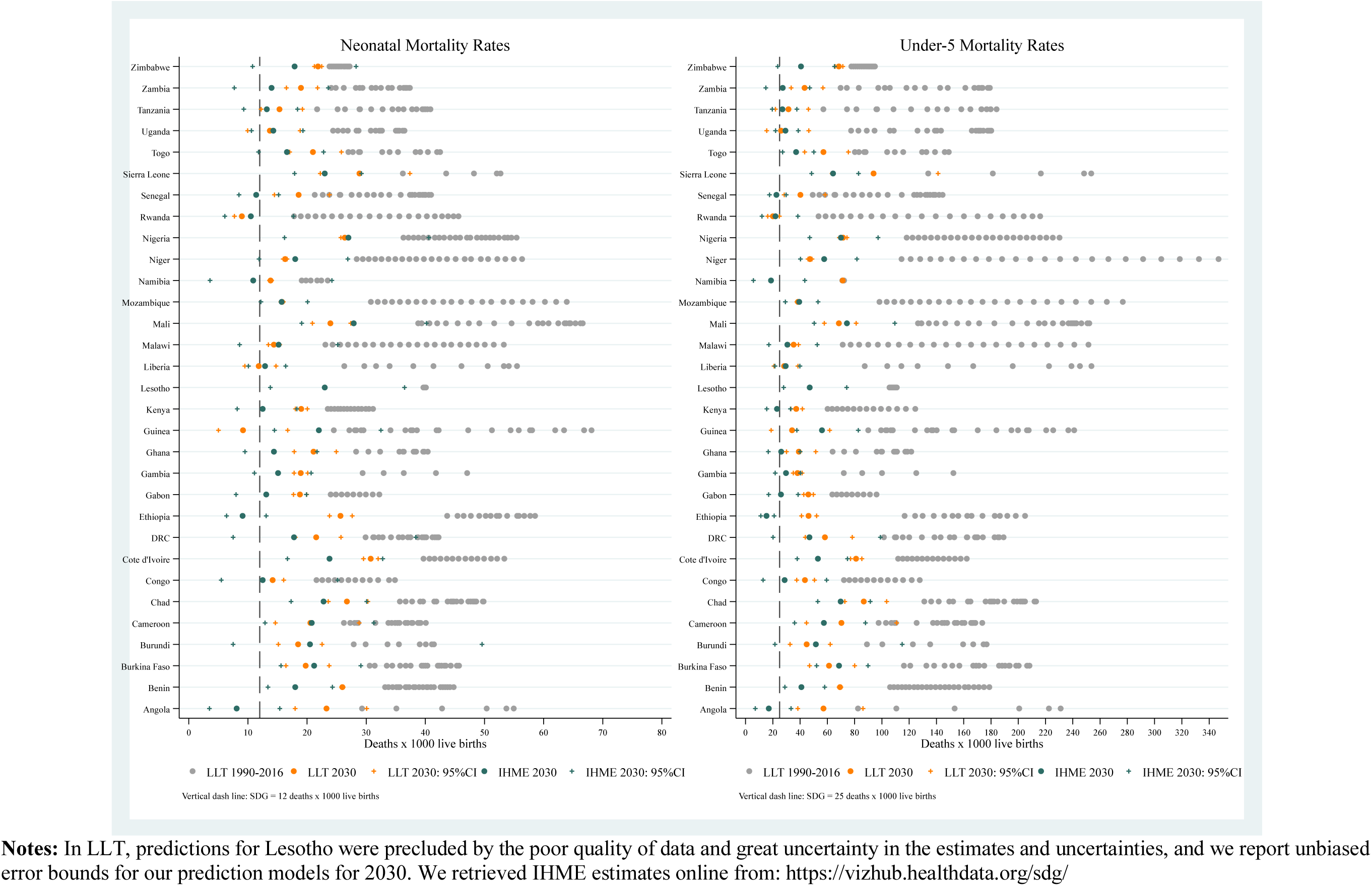
Assessment of the SDG-3 targets for neonatal and under-**5**mortality rates by 2030 based on estimates from the Li-Lee-Tuljapurkar (LLT) model and from the Institute for Health Metrics and Evaluation (IHME) for 31 countries from Sub-Sharan.Africa.

**SI Table.**
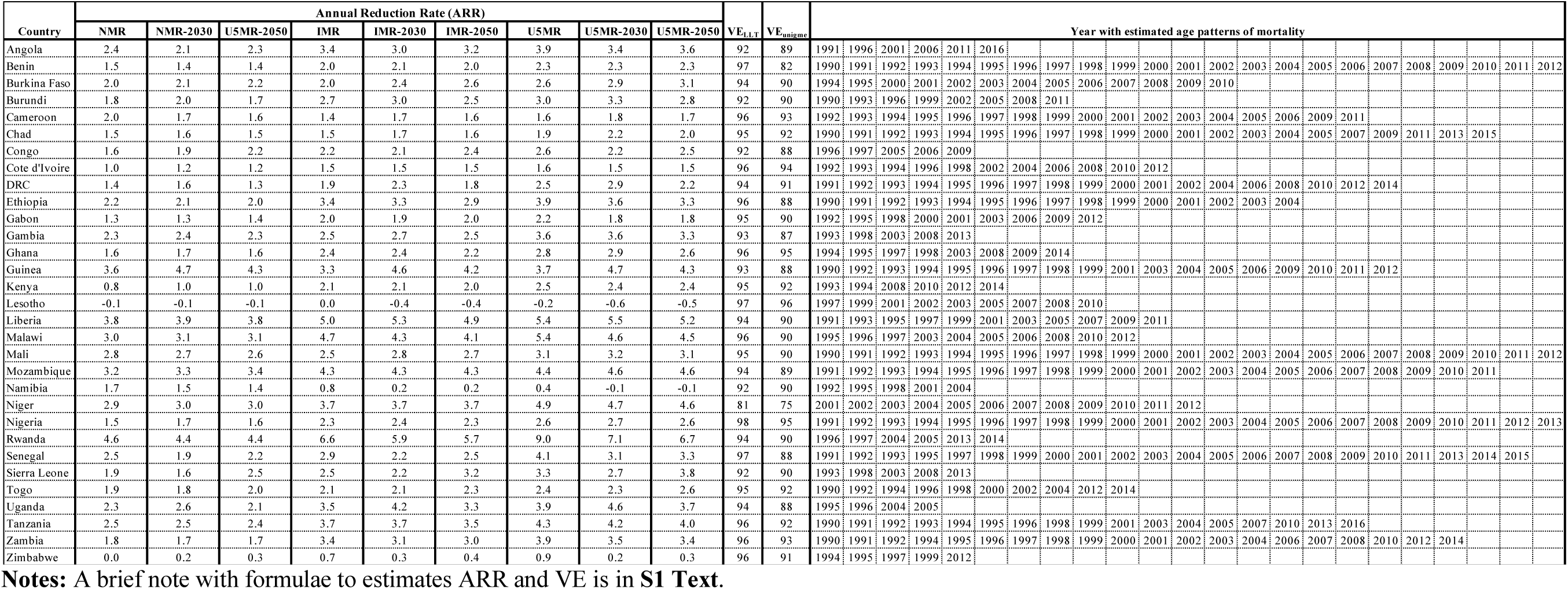
Year of analysis; annual rate of reduction for neonatal (ARR_NMR_), infant (ARR_IMR_), and under-5 mortality rates (ARR_U5MR_); and percentage of variance explained before (VE_LLT_) and after (VE_UNIGME_) adjusting DHS data to match UN IGME estimates

## SI Text. Formal definitions

1) Annual Reduction Rates:

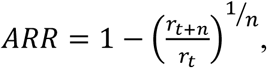

where *r*_*t*_ and *r*_t+n_ are mortality rates at time t and *t*+*n*, respectively, and n is the number of years between *t* and *t*+*n*.

2) Percentage of variance that is explained by the first principal component of the Lee-Carter model:

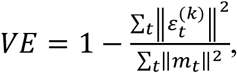

where 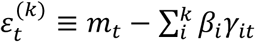 is the error associated with the specification using k principal components. In our LC model, *k*=l1.

## Acknowledgements

Data for this project were accessed using the Stanford Center for Population Health Sciences Data Core. The PHS Data Core is supported by a National Institutes of Health National Center for Advancing Translational Science Clinical and Translational Science Award (UL1 TR001085) and from Internal Stanford funding. The content is solely the responsibility of the authors and does not necessarily represent the official views of the NIH.

